# The mechanism underlying redundant functions of the YTHDF proteins

**DOI:** 10.1101/2022.05.05.490669

**Authors:** Zhongyu Zou, Caraline Sepich-Poore, Xiaoming Zhou, Jiangbo Wei, Chuan He

## Abstract

The YTH *N^6^*-methyladenosine RNA binding proteins (YTHDFs) mediate the functional effects of *N^6^*-methyladenosine (m^6^A) on RNA. Recently, a report proposed that all YTHDFs work redundantly to facilitate RNA decay, raising questions about the exact functions of individual YTHDFs, especially YTHDF1 and YTHDF2. We show that YTHDF1 and YTHDF2 differ in their low-complexity domains (LCDs) and exhibit different behaviors in condensate formation and subsequent physiological functions. Biologically, we also find that the global stabilization of RNA after depletion of all YTHDFs is driven by increased P-body formation and is not strictly m^6^A dependent.

## Background

RNA modification represents an important layer of post-transcriptional regulation of RNA metabolism. Among more than 170 distinct chemical modifications identified on cellular RNA^1^, *N^6^*-methyladenosine (m^6^A) is the most prevalent internal mRNA modification. One of the major pathways through which m^6^A exerts its function is the preferential binding of “reader” proteins to methylated transcripts. Proteins containing the YT521-B homology (YTH) domain, including YTHDF1, 2, and 3 and YTHDC1 and 2 in mammals, are direct m^6^A readers possessing a dedicated m^6^A-binding domain^2–5^. The binding of YTHDF1 to m^6^A-modified mRNAs was shown to induce their translation, which has been linked to various physiologically relevant processes^6–9^. While *Ythdf2* knockout is embryonically lethal in mice^10^, *Ythdf1* knockout mice develop normally within the first three months but exhibit defects in long-term learning and memory^8^. Knockdown of either YTHDF2 or YTHDF3 delays somatic cell reprogramming while YTHDF1 knockdown does not affect this process^11^. These reports suggest that the physiological roles of YTHDF proteins are different, and their functions are specific to certain cellular contexts.

We have reported that the triple knockdown of YTHDF proteins leads to the highest mRNA stabilization compared to the single and double knockdown of YTHDFs^12^. The molecular mechanism behind this observation remains elusive. More recently, one report suggested that YTHDF proteins redundantly function in the decay of methylated RNA and can compensate for each other^13^, and that this redundancy accounts for the significant RNA stabilization after triple knockdown of YTHDFs. The synergistic effect on mRNA decay has also been observed by others with triple knockdown of YTHDF proteins^14,15^; however, various previous reports have also indicated a role for YTHDF1 in translation promotion^16–18^, which made us speculate that a different mechanism may explain the transcriptome stabilization effect observed with YTHDF1-3 triple knockdown.

## Results and discussion

We first examined the notion that YTHDF proteins are not involved in translation regulation, but rather act redundantly to destabilize RNA. This model was presented alongside the concordant idea that YTHDF proteins share highly similar RNA targets, protein partners, and biological functions^13^.

### YTHDF1 promotes translation of its target transcripts

We found two flaws in the analysis of Zaccara *et al*.^13^ that led to the proposal of the redundant model. First, they analyzed the effects of individual YTHDF proteins by grouping RNA by m^6^A modification status, not RNA binding of individual proteins. Specifically, there are 7,105 m^6^ A-modified genes among ~16,000 expressed genes in HeLa cells. Among 6,814 mRNA with translation efficiency data acquired with Ribo-seq, 4,424 m^6^ A-modified mRNA were used by Zaccara *et al*. for their YTHDF1 knockdown analysis (Fig. 1a, left panel), which far exceeds the ~753 high confidence transcripts directly bound by YTHDF1 identified with photoactivatable ribonucleoside-enhanced crosslinking and immunoprecipitation (PAR-CLIP)^3,4,12,13^ (Fig. 1a, right panel). It is not germane to analyze the effects of YTHDF1 knockdown on translation or stabilization of ~60% of the transcriptome as potential targets of YTHDF1 while we know that the protein mainly binds only ~10% of the transcriptome, as many effects on non-targets may be indirect. Our previous m^6^A-QTL studies have already shown that various RBPs can promote or suppress translation through m^6^A^19^. Analyzing all m^6^A-modified mRNAs clearly does not represent the effects of only YTHDF1 or YTHDF2.

**Fig. 1.**
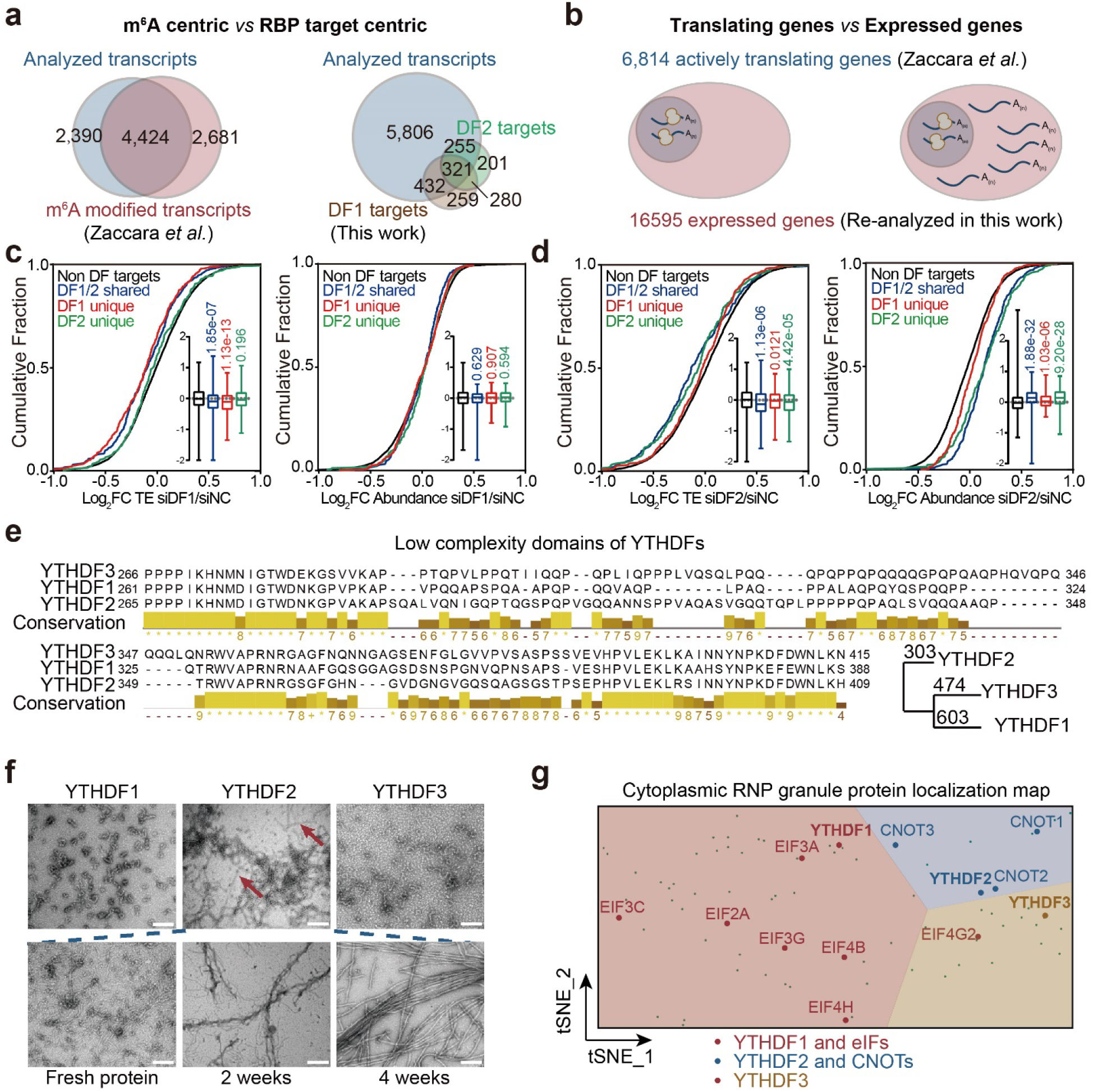
YTHDF1 and YTHDF2 have different molecular functions. (a) Venn diagrams showing RNA analyzed with m^6^A-centric and RBP target-centric analyses. (b) Venn diagrams showing RNA analyzed by Zaccara *et al*. and RNA analyzed in this work. The blue circle denotes translated genes and the red circle denotes expressed genes. (c) Cumulative plots showing changes in RNA translation efficiency (TE, left) and abundance (right) after YTHDF1 knockdown using data from Zaccara *et al*. P values were determined by a Mann-Whitney-Wilcoxon test. (d) Cumulative plots showing changes in RNA translation efficiency (TE, left) and abundance (right) after YTHDF2 knockdown using data from Zaccara *et al*. P values were determined by a Mann-Whitney-Wilcoxon test. (e) Sequence alignment of YTHDF proteins showing differences in the low complexity domain (LCD) region. Conservation scores were calculated by Jalview^31^. (f) Higher-order structure formed by LCDs of YTHDFs under an electron microscopy (EM). Upper panel: fresh protein, lower panel: LCD of YTHDF2 incubated for the indicated times in native buffer. Scale bar: 200 nm. (g) Cytoplasmic RNP granule protein localization map of the cell generated by t-distributed stochastic neighbour embedding (t-SNE) from the CELL MAP project^22^. YTHDF1 or eIFs are delineated in red, YTHDF2 and CNOTs are delineated in blue, and YTHDF3 is delineated in yellow. For boxplots, the center line represents the median, the box limits show the upper and lower quartiles, and whiskers represent 1%-99%. *P* values were determined by a Mann-Whitney-Wilcoxon test.

Second, Zaccara *et al*. analyzed the effects of YTHDFs on mRNA abundance by including only actively translated genes. For their analysis of transcript abundance, the transcripts were pre-filtered to have non-zero read counts in the ribosome-protected fragment (RPF) samples (Fig. 1b). This restrained their analysis to only include actively translated RNA. By studying all detected transcripts (with a sum of > 10 read counts across all samples), we found that the abundance of transcripts with more m^6^A sites actually tends to decrease more upon YTHDF1 or YTHDF3 knockdown (Additional file 1: Fig. S1a). In contrast, only YTHDF2 knockdown and triple knockdown cause stabilization of transcripts with more m^6^A sites (Additional file 1: Fig. S1a). Analyzing only the translated genes, we did not find significant correlations between numbers of m^6^A sites and changes in RNA abundance after YTHDF1 or YTHDF3 knockdown (Additional file 1: Fig. S1a). Interrogating functions of YTHDF proteins only based on actively translated RNA is not appropriate, especially when aiming to elucidate their roles in translation and decay. Moreover, it is inappropriate to draw conclusions about the relative roles of YTHDF proteins in translation and decay based on data from cells treated with a translation inhibitor. Thus, the claim that YTHDF proteins do not affect translation is based on skewed choices of experimental setup and analysis.

To clarify the functional effects of YTHDF proteins on their target mRNA, we grouped transcripts according to their binding by YTHDF1 and YTHDF2 in HeLa cells from published PAR-CLIP datasets^3,4^. Applying these groupings to RNA-seq and ribosome profiling data from knockdown experiments of YTHDF1 and YTHDF2 revealed significant differences. Knockdown of YTHDF1 decreases translation efficiency only of YTHDF1 unique targets and YTHDF1/2 shared targets, while the mRNA abundance of any group of genes is not significantly altered (Fig. 1c). In contrast, YTHDF2 knockdown leads to more significant stabilization of its RNA targets (DF1/2 shared and DF2 unique) (Fig. 1d). We conclude that the major effect of YTHDF1 is to promote the translation of its RNA targets, while YTHDF2 plays a greater role in mRNA stability.

Evolutionarily, *Drosophila* only has one YTHDF ortholog, and it promotes translation of its target mRNA transcripts while having little effect on their abundance^20^. This reinforces the involvement of YTHDF proteins in mRNA translation regulation, not just decay.

### YTHDF1 and YTHDF2 bind different protein partners and form distinct higher-order structures

Zaccara *et al*. also reported that YTHDF proteins bind similar sets of proteins^13^. Amino acid sequences of proteins determine their higher-order structures and molecular functions. If YTHDF proteins share the same protein partners, RNA targets, and biological functions, as was proposed^13^, they should have highly conserved amino acid sequences. However, sequence alignment of human YTHDF proteins shows that they differ greatly in their low-complexity domains, which might lead to distinct features in condensate formation (Fig. 1e). Homology analysis suggests that YTHDF2 is the most different, while YTHDF1 and YTHDF3 are more similar to each other (Fig. 1e, bottom right panel). Indeed, we found that the low-complexity domains (LCDs) of YTHDF2 form fibril-like structures distinct from structures formed by YTHDF1 and YTHDF3 under electron microscopy (Fig. 1f). Elongated incubation facilitates the fibril-like structure formation by YTHDF2 LCDs (Fig. 1f, bottom panel).

The different amino acid sequences and higher-order structures of YTHDF proteins indicate that YTHDF proteins should not share highly similar protein partners. In contrast to the Zaccara *et al*.report^13^, we did not find a shared enrichment of CNOT proteins as high-confidence protein interactors of all three YTHDF proteins by analyzing the same public dataset^21^ (Additional file 1: Fig. S1b). To clarify protein interactions between YTHDFs and either translation or decay machineries, we analyzed a recent protein localization map from HEK293 cells^22^. YTHDF1 tends to display similar subcellular localization in cytosolic RNP granules as eukaryotic initiation factors (eIFs) while YTHDF2 colocalizes better with CNOTs, with YTHDF3 in the margin between CNOTs and eIFs (Fig. 1g and additional file 1: Fig. S1c). These features are in accordance with their reported functions to promote mRNA translation or facilitate decay. Our and others’ results indicate YTHDF1 and YTHDF2 proteins have distinct protein partners and form fundamentally different higher-order structures.

YTHDF2 knockdown or knockout suffices to cause stabilization of its target transcripts, as reported by Zaccara *et al*. and others^23–25^, while YTHDF1 or YTHDF3 single knockdown does not cause significant alteration of RNA abundance. This was attributed to a higher level of YTHDF2 compared to YTHDF1 or YTHDF3 in HeLa cells, by comparing translation efficiency obtained from Ribo-seq^13^. However, we analyzed relative protein levels in HeLa and HEK293T cells and found that the YTHDF1 level is higher in HeLa while YTHDF2 is higher in HEK293T (Additional file 1: Fig. S1d). If YTHDF1 is more abundant than YTHDF2 in HeLa cells, YTHDF2 should not suffice to compensate for YTHDF1 knockdown.

### YTHDF1-3 triple knockdown leads to increased P-body formation and global mRNA stabilization

If each YTHDF protein has a distinct structure and function, how can we explain the synergistic mRNA stabilization observed upon knockdown of all three YTHDF proteins by us, Zaccara *et al*.and others^14^? To answer this, we performed mRNA-seq with spike-in calibration. A slight decrease in mRNA abundance was observed after individual knockdown of each YTHDF protein (Fig. 2a, siDF1, siDF2, siDF3); however, triple knockdown of YTHDF1-3 caused global stabilization of the whole transcriptome (Fig. 2a, siDF1-3). The majority of cytosolic mRNA is stabilized by triple knockdown independent of m^6^A methylation or YTHDF binding (Fig. 2a). Differential gene analyses of datasets obtained from Zaccara *et al*. also show much more significant alteration of the transcriptome (~5-19x more transcripts with adjusted *P* values < 0.05 for differential expression) with YTHDF1-3 triple knockdown than with knockdown of any single YTHDF protein (Additional file 1: Fig. S2a). These results suggest the perturbation of a more fundamental process regulating mRNA stability in the cytosol after YTHDF triple knockdown.

**Fig. 2.**
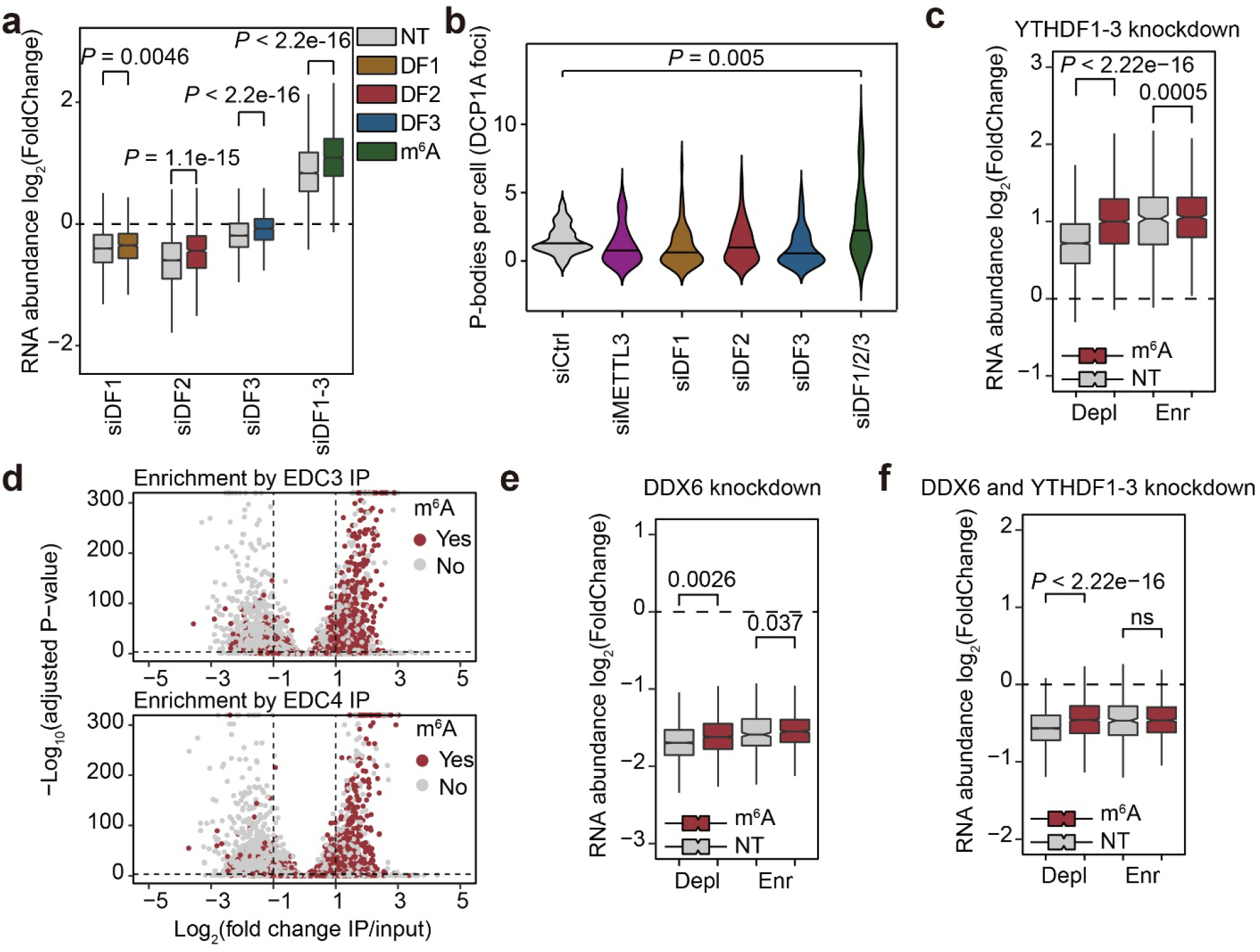
Increased P-body formation accounts for the global RNA stabilization after YTHDF1-3 triple knockdown. (a) Boxplots showing changes in RNA abundance after single knockdown or triple knockdown of YTHDFs in HeLa cells. Grey boxes, non-targets (NT) in each analysis (For siDF1, non-DF1 bound transcripts; for siDF2, non-DF2 bound transcripts; for siDF3, non-DF3 bound transcripts; for siDF1-3, non-methylated transcripts). Colored boxes, YTHDF1 targets (yellow), YTHDF2 targets (red), YTHDF3 targets (blue) and m^6^A modified transcripts (green). (b) Fluorescence microscopy analysis of P-body numbers after single knockdown or triple knockdown of YTHDFs in HeLa cells. Numbers of DCP1A foci per cell were quantified with CellProfiler 3.0. (c) Boxplots showing changes in RNA abundance after triple knockdown of YTHDFs, grouped by both m^6^A modification status and P-body enrichment (depleted: “Depl”, enriched: “Enr”).Red boxes denote m^6^A methylated transcripts and grey boxes denote unmethylated transcripts. (d) Scatter plots showing enrichment of m^6^A-modified transcripts in the P-body enriched pool. Top: EDC3 dataset, bottom: EDC4 dataset. Red dots denote m^6^A methylated transcripts and grey dots denote unmethylated transcripts. (e) Boxplots showing changes in RNA abundance after DDX6 knockdown grouped by both m^6^A modification status and P-body enrichment (depleted: “Depl”, enriched:“Enr”). Red boxes denote m^6^A methylated transcripts and grey boxes denote unmethylated transcripts. (f) Boxplots showing changes in RNA abundance after triple knockdown of YTHDFs in DDX6-depleted HeLa cells grouped by both m^6^A modification status and P-body enrichment (depleted: “Depl”, enriched: “Enr”). Red boxes denote m^6^A methylated transcripts and grey boxes denote unmethylated transcripts. For boxplots, the center line represents the median, the box limits show the upper and lower quartiles, and whiskers represent 1.5 × interquartile range. *P* values were determined by a Mann-Whitney-Wilcoxon test.

We found that depletion of all YTHDF proteins caused increased numbers of processing bodies (P-bodies) in cultured HeLa cells by DCP1A staining (Fig. 2b and additional file 1: Fig. S2b). Cytosolic P-bodies are hubs for RNA processing with reports suggesting roles in facilitating decay or RNA stabilization^26–28^. We decided to characterize the P-body-associated transcriptome in HeLa cells in order to further assess how P-body perturbation may affect m^6^A and non-m^6^A methylated transcripts. We performed RIP-seq using two individual antibodies against EDC3 and EDC4, respectively. The enrichments of transcripts in P-bodies in the two datasets correlate well (Additional file 1: Fig. S2c). Thus, we used the averaged log_2_(enrichment) from these two datasets to define the P-body transcriptome. Unlike P-body depleted (“Depl”) transcripts, m^6^A methylation does not cause a more significant stabilization effect for P-body enriched (“Enr”) transcripts after YTHDF1-3 triple knockdown (Fig. 2c). This indicates that YTHDF1-3 triple knockdown preferentially stabilizes P-body enriched transcripts. We also found that m^6^A-modified transcripts are significantly enriched in the immunoprecipitated fractions in both datasets (Fig. 4d); we categorized transcripts based on numbers of m^6^A peaks and confirmed that groups with more m^6^A peaks are more enriched in P-bodies (Additional file 1: Fig. S2d).

To study whether P-body dynamics account for the transcriptome stabilization observed after YTHDF1-3 triple knockdown, we knocked down DDX6, a protein shown to be essential for P-body assembly^29^. This led to a significant decrease in P-body number visualized after stained with an anti-DCP1A antibody (Additional file 1: Fig. S2e. RNA sequencing with spike-in normalization showed that abolishment of P-bodies leads to global destabilization of RNA (log_2_(FoldChange) < 0) as would be expected (Fig. 2e). Upon YTHDF1-3 triple knockdown, P-body enriched transcripts are more stabilized compared to the P-body depleted transcripts (Fig. 2c). Although P-bodies enrich m^6^A-modified transcripts, our results suggest that the stabilization of transcripts following depletion of all three YTHDF proteins could be a result of increased P-body formation in cells rather than an m^6^A-dependent process, and that the relationship between m^6^A and stabilization after triple knockdown could be confounded by the role and dysregulation of P bodies. Thus, we performed YTHDF protein triple knockdown in DDX6-depleted cells with small interfering RNA (siRNA). The global stabilization effect of YTHDF triple knockdown was completely abolished in DDX6-depleted cells (Fig. 2f). Collectively, our results show that the global stabilization of mRNA after depletion of all YTHDF proteins is a result of increased P-body formation and is not strictly m^6^A dependent.

## Conclusions

The effects of m^6^A on RNA fate are heterogeneous and depend heavily on biological context. The context-dependent roles of m^6^A reader proteins enable the m^6^A-mediated multifaceted regulation of multiple biological processes. YTHDF1 and YTHDF2 proteins have notable sequence differences in their low-complexity regions and form different LLPS granules. Using YTHDF1 target transcripts instead of all methylated mRNA for analysis, we show that YTHDF1 indeed promotes translation in HeLa cells.

In this study, we confirm that depletion of all three YTHDF proteins exhibits a synergistic effect to stabilize mRNA^12–15^. However, we show that this effect is not strictly m^6^A dependent. The triple knockdown of all three YTHDF proteins leads to increased cellular P-body formation and global stabilization of most mRNAs, regardless of their methylation status. Therefore, in addition to the individual functions of these YTHDF proteins that are affected by their relative levels, these abundant proteins also participate in a scaffolding role to maintain cellular RNP granules. Depriving these proteins may expose cellular mRNAs to induce P-body formation for global stabilization of the whole transcriptome.

## Author contributions

Conception, C.H. and Z.Z.; electron microscopy and bacterial protein purification, X.Z.; cell line experiment design and conduction, Z.Z. and J.W.; analysis of public datasets, C.S.-P.; writing, Z.Z., C.S.-P. and J.W.; supervision, C.H..

## Acknowledgements

This work is supported by the National Institute of Health RM1 HG008935 and R01 GM113194 to C.H. C. S.-P. is supported by NIH Medical Scientist Training Program grant T32GM007281 and by NCI F30 CA253987. C.H. is an investigator of the Howard Hughes Medical Institute.

## Materials and methods

### siRNA and plasmid transfections

AllStars negative control siRNA (QIAGEN, 1027281) was used as control siRNA in knockdown experiments. Cells were transfected by using Lipofectamine RNAiMAX (Invitrogen 13778075) for siRNAs (human YTHDF1: QIAGEN SI00764715, human YTHDF2: QIAGEN SI00764757, human YTHDF3: QIAGEN SI04133339, human METTL3: QIAGEN SI05020414 and human DDX6: Dharmacon J-006371-05-0002) according to the manufacturer’s protocols.

### Amino acid sequence alignment

Canonical amino acid sequences of YTHDF1-3 from *Homo sapiens* were retrieved from UniProt Knowledgebase^30^. Sequence alignments were performed with Jalview (version 2.11.2.0)^31^. Conservation scores were calculated with default settings and phylogenetic tree was calculated with the built-in function “Tree” with “Neighbour Joining” and the BLOSUM62 method. Numbers on the tree denote distances in the virtual space depicting similarities between proteins.

### Negative staining transmission electron microscopy

A protein solution (5 μL) of the prion-like domain of each YTHDF was loaded on an EM grid for 10 seconds and excess solution was removed via blotting with filter paper. The grid was then washed with water and stained with 5 μL uranyl acetate (2%) for 15 seconds. All negative staining samples were imaged on a JOEL 1400 microscope.

### Western blot

Protein samples were prepared from respective zebrafish embryos by lysis in RIPA buffer (ThermoFisher Scientific 89900) containing 1 × Halt™ Protease and Phosphatase Inhibitor Cocktail (ThermoFisher Scientific 78441). Protein concentration was measured by NanoDrop 8000 Spectrophotometer (ThermoFisher Scientific). Lysates of equal total protein concentration were heated at 90°C in 1 × loading buffer (Bio-Rad 1610747) for ten minutes. Denatured protein was loaded into 4-12% NuPAGE Bis-Tris gels (Invitrogen NP0335BOX) and transferred to PVDF membranes (ThermoFisher Scientific 88585). Membranes were blocked in Tris-Buffered Saline, 0.1% Tween^®^ 20 (TBST) with 3% BSA (MilliporeSigma A7030) for 30 minutes at room temperature, incubated in a diluted primary antibody solution at 4°C overnight, then washed and incubated in a dilution of secondary antibody conjugated to HRP for 1 hour at room temperature. Protein bands were detected using SuperSignal West Dura Extended Duration Substrate kit (ThermoFisher Scientific 34075) on a FluroChem R (Proteinsimple).

### Fluorescence microscopy

For imaging of P-bodies, HeLa cells were fixed with 4% paraformaldehyde and permeabilized with 0.3% Triton X-100. The blocked coverglass (ThermoFisher Scientific 155409PK) was incubated with an anti-DCP1A-AlexaFluor 488 conjugate (Abcam ab208275) at 4°C overnight. After three washes with DPBS, the nucleus was counterstained with Hoechst 33342 (Abcam ab228551) and the coverglass was kept in DPBS at 4°C before imaging. Samples were imaged on a Leica SP8 laser scanning confocal microscope at the University of Chicago. P-body numbers in each cell were quantified with Cellprofiler 3.0^32^ with a custom workflow.

### RNA-seq library construction and bioinformatic analysis

Library preparation was performed using a SMARTer Stranded Total RNA-Seq Kit v2 (TaKaRa, 634417) following the manufacturer’s protocols. Sequencing was carried out at the University of Chicago Genomics Facility on an Illumina NextSeq machine in single-end mode with 75 base pairs (bp) per read. Raw reads were trimmed with cutadapt (version 1.10)^33^ and then aligned to the human genome and transcriptome (hg38) using HISAT (version 2.1.0)^34^ with the parameter ‘--rna-strandness R’. Annotation files (RefSeq, 2020-04-01, in gtf format) were downloaded from NCBI.

### P-body isolation and analysis

The protocol was adapted from a previous publication^35^. For each immunoprecipitation, HeLa cells from ten 15-cm dishes were collected on ice and flash-frozen with liquid nitrogen before being kept at −80°C until use. The pellet was thawed on ice for 5 min, re-suspended in 1 mL SG lysis buffer (50 mM Tris-HCl pH 7.4, 100 mM KCl, 0.5% NP40, cOmplete mini EDTA-free protease inhibitor (MilliporeSigma 11836170001), 1 U/μl of RNasin Plus RNase Inhibitor (Promega N2611)) and passed through a 25 gauge 5/8 needle attached to a 1 ml syringe 7 times. After lysis, the lysates were spun at 1000 x g for 5 min at 4°C to pellet cell debris. 50 μl and 950 μl of the supernatants were transferred to new microcentrifuge tubes for isolating total and P-body RNAs, respectively. For isolating total RNA, TRIzol™ Reagent (Invitrogen 15596026) was added to the system and RNA was extracted following the manufacturer’s protocol. Following isopropanol precipitation, the RNA pellet was re-suspended in 50 μl RNase-free H_2_O.

The following steps were performed to isolate mammalian P-body cores and extract their RNA: 1) The 950 μl supernatant was spun at 18,000 x g for 20 minutes at 4°C to pellet P-body cores. 2) The resulting supernatant was discarded, and the pellet was re-suspended in 1 ml SG lysis buffer. 3) Steps 1 and 2 were repeated to enrich for SG cores. 4) The resulting pellet was then re-suspended in 300 μl of SG lysis buffer and spun at 850 x g for 2 minutes at 4°C. 5) The supernatant which represents the mammalian SG core enriched fraction was transferred to a new tube. 6) The enriched fraction was pre-cleared twice by adding 60 μL equilibrated DEPC-treated Protein A Dynabeads (ThermoFisher Scientific 10001D) and nutating at 4°C for 30 minutes. Dynabeads were removed using a magnet. 7) 20 μg EDC3 or EDC4 antibody was added to the enriched fraction and nutated at 4°C overnight to affinity purify P-body cores. 8) The solution was spun at 18,000 x g for 20 minutes at 4°C and the supernatant was discarded to remove any unbound antibody. 9) The pellet was then re-suspended in 500 μl SG lysis buffer and 100 μl of equilibrated DEPC-treated Protein A Dynabeads was added. 10) The sample was nutated for 3 hours at 4°C. 11) The Dynabeads were washed three times with wash buffer 1 (20 mM Tris-HCl pH 8.0, 200 mM NaCl, 1 U/μl of RNasin Plus RNase Inhibitor) for 5 minutes, once with wash buffer 2 (20 mM Tris-HCl pH 8.0, 500 mM NaCl, 1 U/μl of RNasin Plus RNase Inhibitor) for 5 minutes, and once with wash buffer 3 (SG lysis buffer + 2 M Urea, 1 U/μl of RNasin Plus RNase Inhibitor) for 2 minutes at 4°C. 12) The beads were resuspended in 200 μl of 100 μg/ml Protease K solution (1X TE buffer, 2M Urea, 1 U/μl of RNasin Plus RNase Inhibitor) and incubated for 15 minutes at 37°C. 13) TRIzol™ Reagent (Invitrogen) was added to the samples and RNA was extracted following the manufacturer’s protocol. Following isopropanol precipitation, the RNA pellet was re-suspended in 20 μl RNase-free H2O. After processing data similarly to the RNA-seq workflow described above through alignment, reads on each NCBI annotated gene were called using the DESeq2 package in R^36^. The fold changes from the DESeq2 output were used as fold enrichment fold in P-bodies.

### RNA seq analysis with spike-in strategy

The same number of cells were counted, and total RNA was isolated with TRIzol™ Reagent (Invitrogen 15596026), according to the manufacturer’s protocol. An amount of ERCC RNA spike-in control (Invitrogen 4456740) proportional to the total cell number was added to each purified total RNA sample before library preparation. After RNA-seq data alignment, reads on each NCBI annotated gene were converted to attomoles by dividing by the sum of reads aligned to the ERCC spike-in. Average log_2_(Fold changes) between attomole amounts were calculated and analyzed.

### Analysis of publicly available RNA-seq data

Raw fastq files were downloaded from Gene Expression Omnibus (accession number GSE134380)^13,37^, and quality checked using FastQC v0.11.5 (http://www.bioinformatics.babraham.ac.uk/projects/fastqc/). Adapters were trimmed using Cutadapt^33^. Reads were then mapped to the hg38 human genome^38^ using HISAT2 v. 2.1.0^39^ with the option --rna-strandedness R. Output sam files were converted to bam files, sorted, and indexed using samtools v. 1.7^40^. Read counts mapping to each gene were obtained using htseq-count^41^ with reference to the hg38 annotation gtf file with options -s reverse, -t exon, -f bam, -i gene_id, -m interserction-nonempty and -r pos. Before further analysis, the sum of counts across all samples was computed for each gene in R v. 4.0.3 using the rowSums function, and genes with 10 or fewer mapped reads were removed. These were taken to represent “expressed” genes. Differential expression analysis of expressed genes was performed with DESeq2^36^ in R v. 4.0.3. Data on *N*^6^-methyladenosine sites and differential expression of “translated” genes in HeLa was obtained directly from processed data files published on Gene Expression Omnibus from the study of interest. Further analyses were performed and plots were created in R v. 4.0.3 using the following packages: biomaRt v. 2.44.4^42^, ggplot2 v. 3.3.5^43^, dplyr v. 1.0.7^44^, ggpubr v. 0.4.0^45^, and forcats v. 0.5.1^46^. When performing analyses involving methylation status, unnamed genes were removed since the published file describing m^6^A sites contained only gene names and m^6^A site numbers. For boxplots, outliers were not shown but were included in statistical analyses.

## References

1 Boccaletto, P. et al. MODOMICS: a database of RNA modification pathways. 2017 update. Nucleic Acids Res 46, D303–D307, doi:10.1093/nar/gkx1030 (2018).

2 Dominissini, D. et al. Topology of the human and mouse m6A RNA methylomes revealed by m6A-seq. Nature 485, 201–206, doi:10.1038/nature11112 (2012).

3 Wang, X. et al. N(6)-methyladenosine Modulates Messenger RNA Translation Efficiency. Cell 161, 1388–1399, doi:10.1016/j.cell.2015.05.014 (2015).

4 Wang, X. et al. N6-methyladenosine-dependent regulation of messenger RNA stability. Nature 505, 117–120, doi:10.1038/nature12730 (2014).

5 Luo, S. & Tong, L. Molecular basis for the recognition of methylated adenines in RNA by the eukaryotic YTH domain. Proc Natl Acad Sci U S A 111, 13834–13839, doi:10.1073/pnas.1412742111 (2014).

6 Han, D. L. et al. Anti-tumour immunity controlled through mRNA m6A methylation and YTHDF1 in dendritic cells (vol 566, pg 270, 2019). Nature 568, E3–E3, doi:10.1038/s41586-019-1046-1 (2019).

7 Weng, Y. L. et al. Epitranscriptomic m(6)A Regulation of Axon Regeneration in the Adult Mammalian Nervous System. Neuron 97, 313–325 e316, doi:10.1016/j.neuron.2017.12.036 (2018).

8 Shi, H. L. et al. m(6)A facilitates hippocampus-dependent learning and memory through YTHDF1. Nature 563, 249–+, doi:10.1038/s41586-018-0666-1 (2018).

9 Gokhale, N. S. et al. N6-Methyladenosine in Flaviviridae Viral RNA Genomes Regulates Infection. Cell Host Microbe 20, 654–665, doi:10.1016/j.chom.2016.09.015 (2016).

10 Zhao, B. S. et al. m(6)A-dependent maternal mRNA clearance facilitates zebrafish maternal-to-zygotic transition. Nature 542, 475–478, doi:10.1038/nature21355 (2017).

11 Liu, J. et al. YTHDF2/3 Are Required for Somatic Reprogramming through Different RNA Deadenylation Pathways. Cell Rep 32, 108120, doi:10.1016/j.celrep.2020.108120 (2020).

12 Shi, H. et al. YTHDF3 facilitates translation and decay of N(6)-methyladenosine-modified RNA. Cell Res 27, 315–328, doi:10.1038/cr.2017.15 (2017).

13 Zaccara, S. & Jaffrey, S. R. A Unified Model for the Function of YTHDF Proteins in Regulating m(6)A-Modified mRNA. Cell 181, 1582–1595 e1518, doi:10.1016/j.cell.2020.05.012 (2020).

14 Lasman, L. et al. Context-dependent functional compensation between Ythdf m(6)A reader proteins. Genes Dev 34, 1373–1391, doi:10.1101/gad.340695.120 (2020).

15 Kontur, C., Jeong, M., Cifuentes, D. & Giraldez, A. J. Ythdf m(6)A Readers Function Redundantly during Zebrafish Development. Cell Rep 33, 108598, doi:10.1016/j.celrep.2020.108598 (2020).

16 Liu, X. et al. YTHDF1 Facilitates the Progression of Hepatocellular Carcinoma by Promoting FZD5 mRNA Translation in an m6A-Dependent Manner. Mol Ther Nucleic Acids 22, 750–765, doi:10.1016/j.omtn.2020.09.036 (2020).

17 Pi, J. et al. YTHDF1 Promotes Gastric Carcinogenesis by Controlling Translation of FZD7. Cancer Res 81,2651–2665, doi:10.1158/0008-5472.CAN-20-0066 (2021).

18 Jin, H. et al. N(6)-methyladenosine modification of ITGA6 mRNA promotes the development and progression of bladder cancer. EBioMedicine 47, 195–207, doi:10.1016/j.ebiom.2019.07.068 (2019).

19 Zhang, Z. et al. Genetic analyses support the contribution of mRNA N(6)-methyladenosine (m(6)A) modification to human disease heritability. Nat Genet 52, 939–949, doi:10.1038/s41588-020-0644-z (2020).

20 Kan, L. et al. A neural m(6)A/Ythdf pathway is required for learning and memory in Drosophila. Nat Commun 12, 1458, doi:10.1038/s41467-021-21537-1 (2021).

21 Youn, J. Y. et al. High-Density Proximity Mapping Reveals the Subcellular Organization of mRNA-Associated Granules and Bodies. Mol Cell 69, 517–532 e511, doi:10.1016/j.molcel.2017.12.020 (2018).

22 Go, C. D. et al. A proximity-dependent biotinylation map of a human cell. Nature 595, 120–124, doi:10.1038/s41586-021-03592-2 (2021).

23 Fang, R. et al. EGFR/SRC/ERK-stabilized YTHDF2 promotes cholesterol dysregulation and invasive growth of glioblastoma. Nat Commun 12, 177, doi:10.1038/s41467-020-20379-7 (2021).

24 Heck, A. M., Russo, J., Wilusz, J., Nishimura, E. O. & Wilusz, C. J. YTHDF2 destabilizes m(6)A-modified neural-specific RNAs to restrain differentiation in induced pluripotent stem cells. RNA 26, 739–755, doi:10.1261/rna.073502.119 (2020).

25 Lee, Y., Choe, J., Park, O. H. & Kim, Y. K. Molecular Mechanisms Driving mRNA Degradation by m(6)A Modification. Trends Genet 36, 177–188, doi:10.1016/j.tig.2019.12.007 (2020).

26 Hubstenberger, A. et al. P-Body Purification Reveals the Condensation of Repressed mRNA Regulons. Mol Cell 68, 144–157 e145, doi:10.1016/j.molcel.2017.09.003 (2017).

27 Luo, Y., Na, Z. & Slavoff, S. A. P-Bodies: Composition, Properties, and Functions. Biochemistry 57, 2424–2431, doi:10.1021/acs.biochem.7b01162 (2018).

28 Wang, C. et al. Context-dependent deposition and regulation of mRNAs in P-bodies. Elife 7, doi:10.7554/eLife.29815 (2018).

29 Ayache, J. et al. P-body assembly requires DDX6 repression complexes rather than decay or Ataxin2/2L complexes. Mol Biol Cell 26, 2579–2595, doi:10.1091/mbc.E15-03-0136 (2015).

30 UniProt, C. UniProt: the universal protein knowledgebase in 2021. Nucleic Acids Res 49, D480–D489, doi:10.1093/nar/gkaa1100 (2021).

31 Waterhouse, A. M., Procter, J. B., Martin, D. M., Clamp, M. & Barton, G. J. Jalview Version 2--a multiple sequence alignment editor and analysis workbench. Bioinformatics 25, 1189–1191, doi:10.1093/bioinformatics/btp033 (2009).

32 McQuin, C. et al. CellProfiler 3.0: Next-generation image processing for biology. PLoS Biol 16, e2005970, doi:10.1371/journal.pbio.2005970 (2018).

33 Martin, M. Cutadapt removes adapter sequences from high-throughput sequencing reads. 2011 17, 3, doi:10.14806/ej.17.1.200 (2011).

34 Kim, D., Langmead, B. & Salzberg, S. L. HISAT: a fast spliced aligner with low memory requirements. Nat Methods 12, 357–360, doi:10.1038/nmeth.3317 (2015).

35 Khong, A. et al. The Stress Granule Transcriptome Reveals Principles of mRNA Accumulation in Stress Granules. Mol Cell 68, 808–820 e805, doi:10.1016/j.molcel.2017.10.015 (2017).

36 Love, M. I., Huber, W. & Anders, S. Moderated estimation of fold change and dispersion for RNA-seq data with DESeq2. Genome Biol 15, 550, doi:10.1186/s13059-014-0550-8 (2014).

37 Edgar, R., Domrachev, M. & Lash, A. E. Gene Expression Omnibus: NCBI gene expression and hybridization array data repository. Nucleic Acids Res 30, 207–210, doi:10.1093/nar/30.1.207 (2002).

38 Frankish, A. et al. GENCODE reference annotation for the human and mouse genomes. Nucleic Acids Res 47, D766–D773, doi:10.1093/nar/gky955 (2019).

39 Kim, D., Paggi, J. M., Park, C., Bennett, C. & Salzberg, S. L. Graph-based genome alignment and genotyping with HISAT2 and HISAT-genotype. Nat Biotechnol 37, 907–915, doi:10.1038/s41587-019-0201-4 (2019).

40 Li, H. et al. The Sequence Alignment/Map format and SAMtools. Bioinformatics 25, 2078–2079, doi:10.1093/bioinformatics/btp352 (2009).

41 Anders, S., Pyl, P. T. & Huber, W. HTSeq--a Python framework to work with high-throughput sequencing data. Bioinformatics 31, 166–169, doi:10.1093/bioinformatics/btu638 (2015).

42 Durinck, S., Spellman, P. T., Birney, E. & Huber, W. Mapping identifiers for the integration of genomic datasets with the R/Bioconductor package biomaRt. Nat Protoc 4, 1184–1191, doi:10.1038/nprot.2009.97 (2009).

43 Wickham, H. D. N., and Thomas Lin Peterson. ggplot2: elegant graphics for data analysis. (springer, 2016).

44 Wickham H, F. R., Henry L, Müller K dplyr: A Grammar of Data Manipulation., <https://dplyr.tidyverse.org, https://github.com/tidyverse/dplyr> (2022).

45 Kassambara, A., and Maintainer Alboukadel Kassambara.. “Package ‘ggpubr’.”R package version 0.4.4 6 (2020).

46 Wickham, H. “Forcats: Tools for Working with Categorical Variables (Factors); R package verion 0.5. 1; 2021.”. (2021).

