## Supplemental figures for "The mechanism underlying redundant functions of the YTHDF proteins"

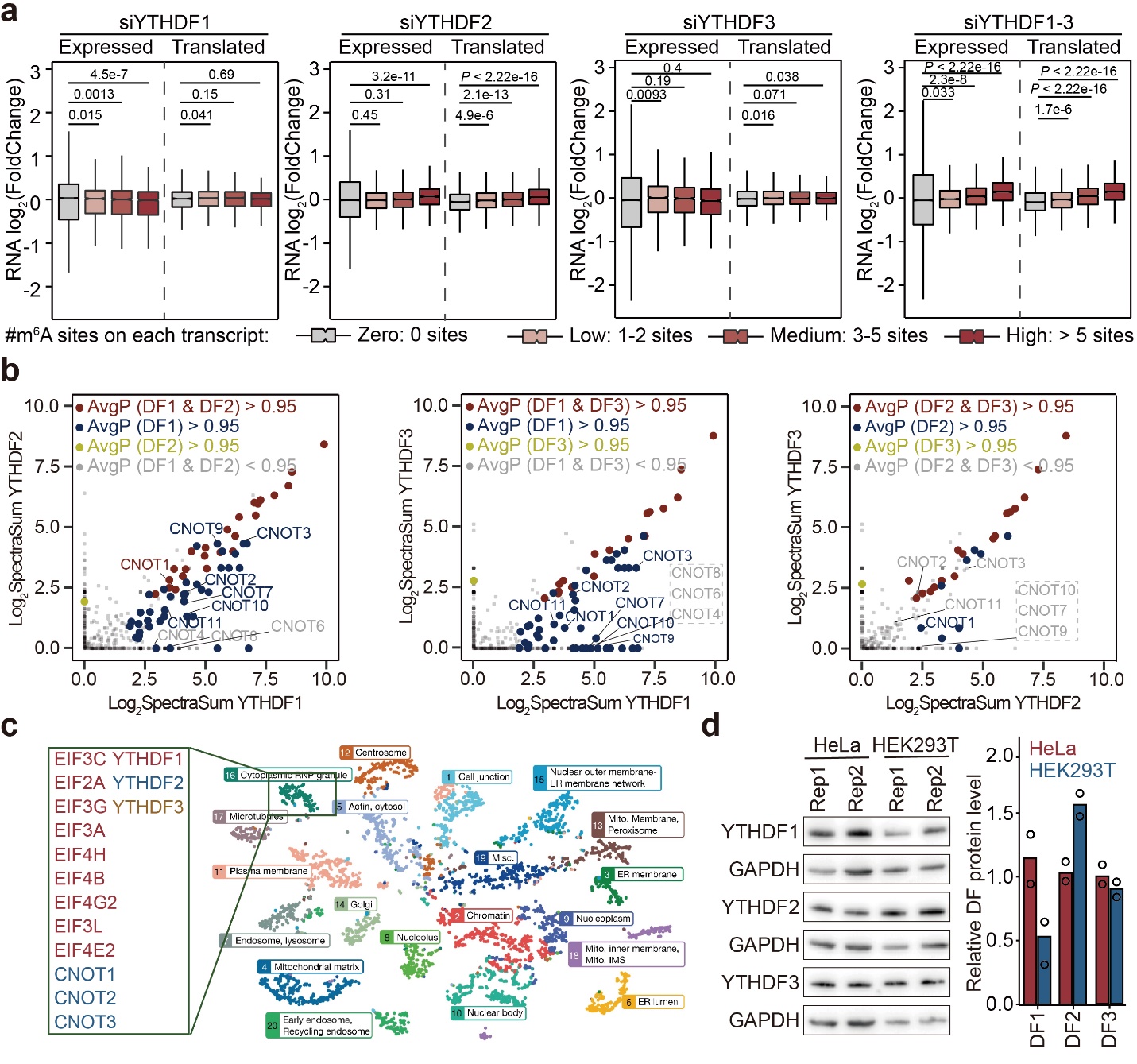

**Fig. S1 Discrepancies between experimental results and the “unified” model of YTHDFs**

(a) Boxplots showing fold changes of transcripts with different numbers of m^6^A sites when analyzing all expressed genes (“Expressed”) or only translated genes (“Translated”) after individual knockdown of YTHDFs. Transcripts were classified by individual numbers of m^6^A sites (zero: 0 sites, low: 1-2 sites, medium: 3-5 sites, high: more than 5 sites). For boxplots, the center line represents the median, the box limits show the upper and lower quartiles, whiskers represent 1.5 × interquartile range. *P* values were determined by a Mann-Whitney-Wilcoxon test.
(b) Scatter plots showing comparisons of YTHDF protein interactions between YTHDF1/YTHDF2 (left), YTHDF1/ YTHDF3 (middle), and YTHDF2/YTHDF3 (right). Processed Bio-ID results were obtained from Youn *et al.^1^*.
(c) Demonstration of the original protein localization map of the cell generated by t-distributed stochastic neighbour embedding (t-SNE) from the CELL MAP project^2^. The cluster annotated as cytoplasmic RNP granules containing YTHDF proteins was selected for analysis in figure 1g.
(d) Western blots showing relative protein levels of YTHDFs in WT HeLa and HEK293T cells. Right: quantification of western blots using ImageJ.

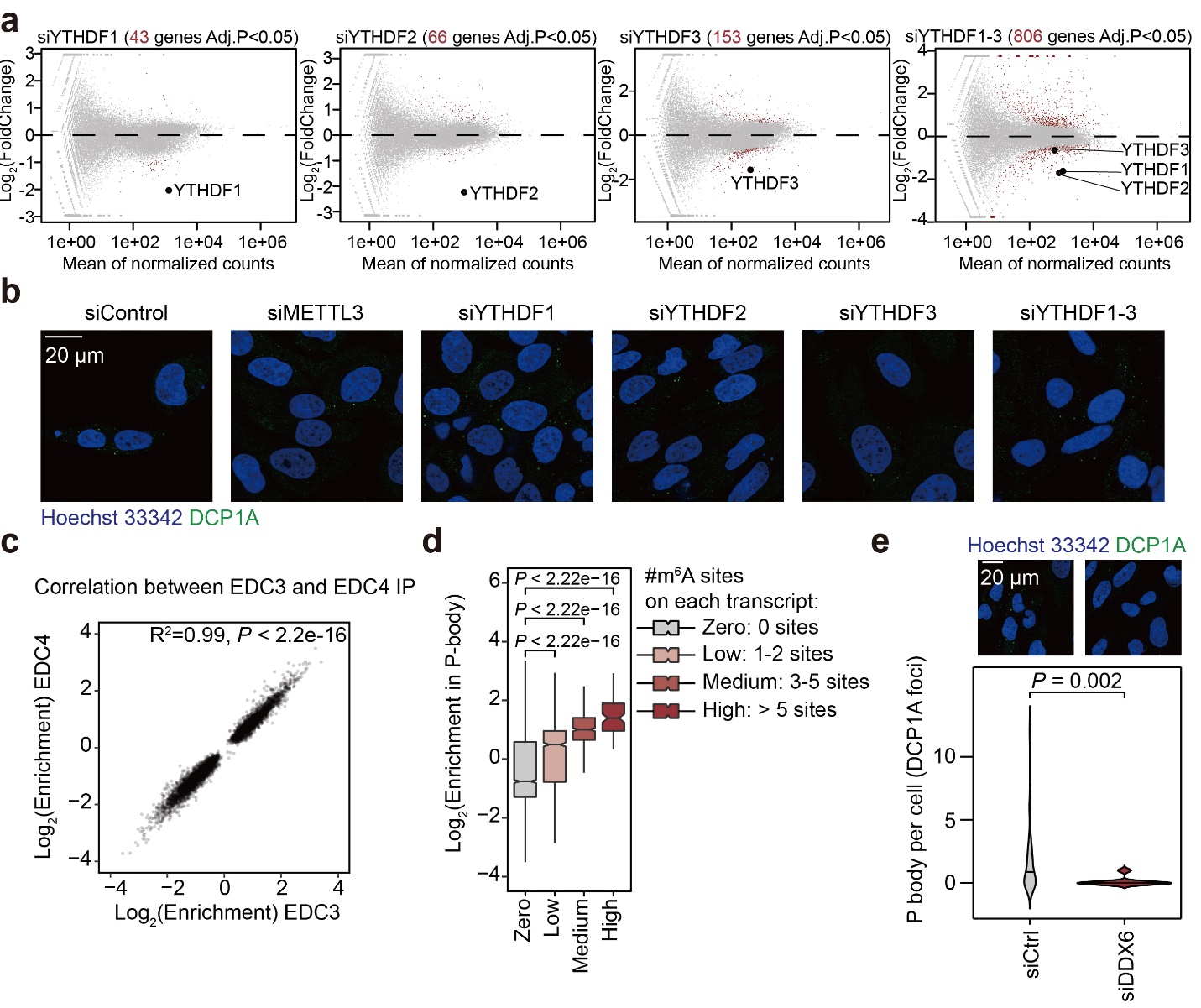

**Fig. S2 Increased P-body formation after YTHDF1-3 triple knockdown.**

(a) MA plots showing RNA expression changes after single knockdown or triple knockdown of YTHDFs. Grey dots denote genes not significantly altered and red dots denote significantly differentially expressed genes. Genes targeted by siRNAs are labeled.
(b) Representative images from P-body imaging in HeLa cells, related to figure 2b. P-bodies were stained using a DCP1A antibody and cell nuclei were counterstained by Hoechst 33342. Eight images were captured for each condition in n = 3 individual experiments.
(c) Scatter plot showing correlation between RNA enrichments in P-body transcriptomes determined with EDC3 or EDC4 antibodies. *P* value was determined by Pearson’s correlation.
(d) Boxplots showing enrichments of different transcripts grouped by numbers of m^6^A sites identified by MeRIP-seq (zero: 0 sites, low: 1-2 sites, medium: 3-5 sites, high: more than 5 sites). For boxplots, the center line represents the median, the box limits show the upper and lower quartiles, whiskers represent 1.5 × interquartile range. *P* values were determined by a Mann-Whitney-Wilcoxon test.
(e) Fluorescence microscopy analysis of P-body numbers after knockdown of DDX6 in HeLa cells. Numbers of DCP1A foci per cell were quantified with CellProfiler 3.0. *P* values were determined by a Mann-Whitney-Wilcoxon test.

**References**

1	Youn, J. Y. *et al.* High-Density Proximity Mapping Reveals the Subcellular Organization of mRNA-Associated Granules and Bodies. *Mol Cell* **69**, 517-532 e511, doi:10.1016/j.molcel.2017.12.020 (2018).

2	Go, C. D. *et al.* A proximity-dependent biotinylation map of a human cell. *Nature* **595**, 120-124, doi:10.1038/s41586-021-03592-2 (2021).
